# Finite-state discrete-time Markov chain models of gene regulatory networks

**DOI:** 10.1101/006361

**Authors:** V.P. Skornyakov, M.V. Skornyakova, A.V. Shurygina, P.V. Skornyakov

## Abstract

In this study Markov chain models of gene regulatory networks (GRN) are developed. These models gives the ability to apply the well known theory and tools of Markov chains to GRN analysis. We introduce a new kind of the finite graph of the interactions called the combinatorial net that formally represent a GRN and the transition graphs constructed from interaction graphs. System dynamics are defined as a random walk on the transition graph that is some Markovian chain. A novel concurrent updating scheme (evolution rule) is developed to determine transitions in a transition graph. Our scheme is based on the firing of a random set of non-steady state vertices of a combinatorial net. We demonstrate that this novel scheme gives an advance in the modeling of the asynchronicity. Also we proof the theorem that the combinatorial nets with this updating scheme can asynchronously compute a maximal independent sets of graphs. As proof of concept, we present here a number of simple combinatorial models: a discrete model of auto-repression, a bi-stable switch, the Elowitz repressilator, a self-activation and show that this models exhibit well known properties.

## 1 Introduction

Efforts to study gene expression regulation networks has led to a detailed description many of them, and many more are to be identified in the near future. Therefore there is a need to develop methods of computational and theoretical analysis of GRNs. One of the most promising directions is to reduce the problem to the study of Markov chains, generated in some way from the GRN [1, 2, 3, 4, 5]. Usually Boolean networks [6] are used as a formal representation of GRN. Classification of process states, the study of long-term behavior [7], and development of optimal strategies for therapeutic intervention [8, 9, 10, 7, 11, 12, 13, 14, 15] provide good examples of this approach [16]. In contrast to the Boolean network, the Hopfield networks are defined using arithmetic operations [17]. It is a well-developed branch of science which deals with stochastic processes of asynchronous state switching as a result of interactions. As such they are similar to Boolean networks. A Hopfield like formalism also leads to the definition of the Markov chain. In the Hopfield network area essential results were obtained in the study of various update schemes [18], network oscillations [19], solving of combinatorial optimization problems [20, 21, 22, 23, 19, 24, 25, 26], and estimating the rate of the convergence and many others. This makes it valuable to study the possibility of using Hopfield like networks for the construction of Markov chains from GRNs and other interactions graphs. We consider a GRN as a kind of interaction graph. Interaction (regulatory) graphs are emerged in various fields of the life science [27]. Nowadays, their transition graphs are often used to analyze properties of interactions (regulations). One promising way to understand the nature of the regulation or interactions represented by interaction graphs is to analyze the Markovian chain associated with their transition graphs.

## 2 Method

The proposed method may be viewed as a version of the Hopfield like network [17] where groups of randomly selected *unstable units* are updated in parallel [18].

### 2.1 The interactions graph and non steady state vertices

Let *G* = (*V*, *E*) be a directed graph, where *V* is set of vertices and *E* is set of edges. Let *B* = {0, 1} be a set of vertex states. We say that the vertex is *active* if the state of this vertex is equal to “1”, otherwise we say that vertex is *inactive*. The map function *M*: *V* → *B* give us the state of each vertex. If some vertex *υ* ∈ *V*, then *M*(*υ*) is the state of the vertex *υ* that correspond to map *M*. *M*(*υ*) = 1 is equivalent to the vertex v is active under map M. *M*(*υ*) = 0 is equivalent to the vertex v is inactive. The weight function *W*: *E* → *R* gives the value of each edge of the graph *G*, which represent the power of interactions. If *e* = (*u*, *υ*) than we say that *u* is a direct ancestor of *υ* and *υ* is a direct descendant of *u*. The *influence* on *υ under* the map *M* is defined as the sum of weights of edges from all direct active ancestors of vertex *υ*. The influence on *υ* under the map *M* is denoted by *I*(*υ*, *M*) (also called “the local field or the net input”). That is

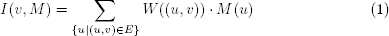

This influence is determined by map function *M* and weight function *W*. Only if the weight function *W* is assumed to be constant over time then we say that influence *I*(*υ*, *M*) is influence of map *M* on vertex *υ*. If *I*(*υ*, *M*) ≥ 0 we say that map *M* activate vertex *υ* otherwise we say that map *M* repress vertex *υ*. Now we can give most important definition of vertex steady state under the map *M*. Let *υ* be the active under map M, if map *M* activate *υ* then the state of *υ* is a steady state under the map *M*, else the state of *υ* is a non steady state under the map *M*. Let *υ* be inactive under map *M*, if map *M* inactivate *υ* then the state of *υ* is a steady state under the map *M*, else the state of *υ* is a non steady state under the map *M*. If all vertices are in steady state under the map M we say that map M is steady map. The *forced* state of the vertex *υ* under map *M* is defined as follows:

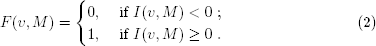

By definition, if forced state and current state of *υ* are the same then state of the vertex *υ* under the map M is steady. Next equation give us the formal definition: 
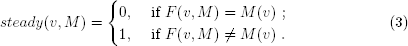

### 2.2 The Random Set update rule

Now we consider a stochastic process{*Y_j_*, *j* = 1, 2, 3,…} that takes on the set of maps of some interaction graph *G*, where *Y_j_* denote the map of *G* at time period *j*. At each time period *j* for each non steady-state vertex *υ_i_* under the map *Y_j_* we change the current vertex state to a forced state with the probability *p_i_*, and the current state stays unchanged with the probability 1 − *p_i_*. Let *S* = {*υ*_1_, *υ*_2_, …, *υ_n_*} be a set of all non steady-state vertices at the time period *j*. Vertices chosen to change their state in one-step transition form a random set *X* ⊆ *S*. To make the new map that direct accessible from current map *Y_j_* all vertices from *X* simultaneously change their state whereas another vertices stay unchanged. Let P= {*p*_1_, *p*_2_, …, *p_n_*} be some vector of numbers such that 0 ≤ *p_i_* ≤ 1, we refer to *p_i_* as the probability of the state changing (firing) of the non steady-state vertex *υ_i_*. For any *X* ⊆ *S* let 1_*X*_: *S* → *B* be indicator function such that 1_*X*_(*υ_i_*) = 1 if *υ_i_* ∈ *X*, otherwise 1_*X*_(*υ_i_*) = 0. Let **P**_*X*_: *S* → [0.0, 1.0] denote the function such that **P**_*X*_(*υ_i_*) = *p_*i*_* if 1_*X*_(*υ*_*i*_) = 1, otherwise **P**_*X*_(*υ_i_*) = 1 − *p_i_*. Hence we assume that each *υ_i_* acts independently to make random set *X* then the production of **P**_*X*_ give us *p_X_* that is the firing probability of the random set *X*

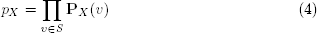

Evidently,

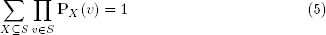

Now we can apply this definition of Random Set update rule and its probabilities to define the transitions graph of the combinatorial net models.

### 2.3 The transitions graph of the interactions graph

Let *S* be the set of non steady state vertices under map *M* = *Y_j_*. *S* represent a full set of vertices each of which can flip to a forced state at next *j*+1 time step. In the combinatorial model steady state vertices can not flip. To construct a transition graph we should define the full set of maps *M*_1_, *M*_2_, … that are direct reachable from the map *M*. Each pair (*M*, *M_i_*) will correspond to one edge in transition graph. What is the set of maps *M*_1_, *M*_2_, …, which can be direct (by one-step transition) reached from map *M*? To represent the independence and the asynchronicity we assume that any random set of non steady state vertices *X* ⊆ *S* can produce next map from current map. Let *M_X_* be map such that *M_X_*(*υ*) = *F*(*υ*, *M*)) if *υ* ∈ *X* and *M_X_*(*υ*) = *M*(*υ*) if *υ* ∉ *X*, then we say that *M_X_* produced by *X* from *M*. That is,the random set *X* of non steady state vertices produce the map *M_X_* from a map *M*. The weight of edge from map *M* to a map *M_X_* are given by the probability defined by equation (4).

### 2.4 The random walk network dynamics

We suppose that whenever the process is in state *M*, there is a probability *p_X_* such that at next step it will be in state *M_X_*. This probability is defined for each random set *X* of non-steady state vertices of map *M*.

### 3 Random Set update rules are more general

It is well known that asynchronous and Random Set update rules are equivalent in the sense of global stable states [28]. But in the sense of the reachability of one state from another they are not equivalent. Figure 1 show the Mace combinatorial model that illustrate this fact. The vertex *e* provides constant level of repression for vertex *c*, that is equal −2. Let the vertex *d* of the Mace will be active at the start. Then it can activate both middle vertices *a* and *b*. Due to repression, the vertex *c* of the Mace can be activated only if both middle vertices *a* and *b* will be active simultaneously. Asynchronous (one at a time) updating exclude simultaneous activation of these vertices, but Random Set update rule do not. Synchronous update rule do not exclude simultaneous activation of a and *b*, but it make the system deterministic. The Random Set update rule is more general than both synchronous and asynchronous update rules, because it allow all possible paths of a system evolution. Therefore, transition graphs of both synchronous and asynchronous update rules are subgraphs of Random Set update graph.

**Figure 1:**
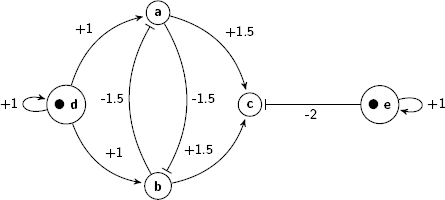
The Mace

## 4 Examples of combinatorial models

Next we use described above method to develop models of some important graphs of repressive interactions of self activating nodes and prove their main properties. Such models we call combinatorial models.

### 4.1 The combinatorial model of an Auto-repression

A negative auto-regulation or an auto-repression occurs when products of some gene represses its own gene. This form of a simple regulation serve as a basic building block of most important transcription networks [27, 29]. An Auto-repression can produce oscillations. For example embryonic stem cells fluctuate between high and low Nanog expression and the Nanog activity is auto-repressive [30]. Our model of an auto-repression shown in Figure 3a and Figure 2a also exhibit stochastic oscillating behavior. Figure 2a present a graph of interactions G = (V, E) of the Auto-repression model. The set V contains a single vertex *υ*_1_. And the set E consist of a single edge *e*1 = (*υ*1, *υ*1) from vertex v1 to itself. The weight of e1 is equal to −1. Let the *M*_0_ be a starting map and *M*_0_(*υ*_1_) = 0, i.e. *υ*_1_ is inactive under the *M*_0_. Therefore, influence of *M*_0_ to vertex *υ*_1_ equals −1, *I*(*υ*1, *M_0_*) = −1. By equation [2], *F*(*υ*_1_, *M*_0_) = 0, then state of the vertex *υ*_1_ is non steady under the map *M*_0_. Let *p*_1_ = 1/2 then the state of the vertex *υ*_1_ could be changed to active state with probability 1/2 and could stay unchanged with probability 1/2. The another state of the model are also non steady. Therefore, there is only non steady states in the model. Thus, it will oscillate infinitely between 0 and 1. Figure 3a show full transition graph of the auto-repression model.

**Figure 2:**
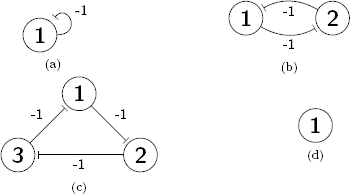
The interaction graph of: (a) The Autorepression model; (b) The Bistable Switch model; (c) The Elowitz repressilator model; (d) The self-activation model.

**Figure 3:**
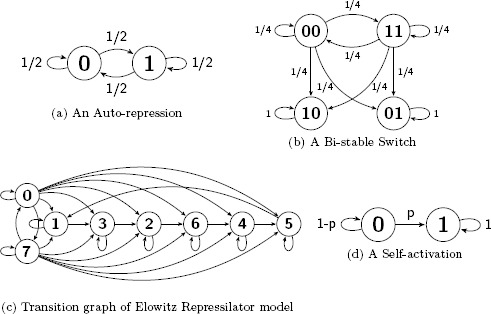
The Transition Graphs of: (a) The Autorepression model; (b) The Bistable Switch model; (c) The Elowitz repressilator model; (d) The self-activation model.

### 4.2 The combinatorial model of a Bi-stable Switch

A Bi-stable Switch is bi-stable gene regulatory network that constructed from a two mutually repressive genes [31]. They are widely common in nature and most used in synthetic biology [32, 33]. ODEs used to construct their mathematical models are convenient way for analyzing in detail some small circuits. Here we develop techniques that can be used to construct models of large networks of bis-table switches and to prove some their important properties. Thus, we use probabilistic coarse-scale modeling approach [34] instead of fine-scale ODE modeling. Our model of Bis-table Switch shown in Fig 2b and Fig 3b exhibit two steady states. Figure 2b present a graph of interactions *G* = (*V*, *E*) of the Bi-stable Switch model. The set *V* = {*υ*_1_, *υ*_2_} contains two vertices and *E* contains two edges, which weights are −1. Probabilities of firing non steady state vertices let be 1/2. Figure 3b present the transition graph of the model. Maps 00 and 11 are non steady, whereas maps 01 and 10 are steady. To show that a map *M* is steady we must show that each vertex *υ* is steady under map *M*. To show that a vertex *υ* is steady under map *M* we must compute value of influence *F*(*υ*, *M*) of map *M* on vertex *υ* and compare it to the state *υ* under map *M*, i.e. *M*(*υ*). Now we will show that map 01 is steady.

### 4.3 The combinatorial model of the Elowitz Repressilator

Elowitz repressilator consists of three genes [35]. Each of these genes are constitutively expressed. The first gene inhibits the transcription of the second gene, whose protein product in turn inhibits the expression of a third gene, and finally, third gene inhibits first gene expression, completing the cycle. Such a negative feedback loop lead to oscillations. The combinatorial model of Elowitz repressilator produce oscillations and consists of three vertices and three edge, which weights is equal to −1. Figure 2c present a graph of interactions *G* = (*V*, *E*) of the Elowitz repressilator model, where the set *V* = {*υ*_1_, *υ*_2_, *υ*_3_} contains three vertices and *E* = {(*υ*_1_, *υ*_2_), (*υ*_2_, *υ*_3_), (*υ*_3_, *υ*_1_)} contains three edges. Oscillations are produced by circle {1,3,2,6,4,5} in Figure 3c. Decimal, binary, and graphical representations of the state space of the Elowitz repressilator are shown in Table 1.

**Table 1:**
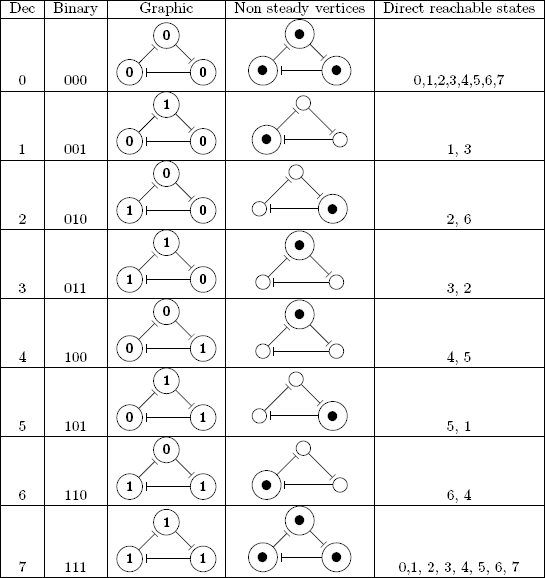
Decimal, binary, and graphical representations of the state space of the Elowitz repressilator

| Dec | Binary | Graphic | Non steady vertices | Direct reachable states |
| --- | --- | --- | --- | --- |
| 0 | 000 |  |  | 0,1,2,3,4,5,6,7 |
| 1 | 001 |  |  | 1, 3 |
| 2 | 010 |  |  | 2, 6 |
| 3 | 011 |  |  | 3, 2 |
| 4 | 100 |  |  | 4, 5 |
| 5 | 101 |  |  | 5, 1 |
| 6 | 110 |  |  | 6, 4 |
| 7 | 111 |  |  | 0,1, 2, 3, 4, 5, 6, 7 |

### 4.4 The combinatorial model of the self-activation

A constitutively expressed gene represent an example of a self-activation. Such gene do not require any interaction to be active. A combinatorial model of the self-activation consist of one vertex and no edges. In any case the influence on it equals 0 since there are no other vertices. Therefore a forced state of the vertex equals 1. Thus 1 is steady state and 0 is non steady state. Vertex starting in steady state will stay in it infinitely. Vertex starting in non steady state with probability p flip to steady state and with probability 1-p stay in non steady state. The amount of time periods *T* which the vertex spent in non steady state is the random variable. The distribution of this random variable is the shifted geometric distribution with parameter p. Figures 2d and 3d represents the combinatorial model of a self-activation.

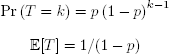

### 5 The network of bi-stable switches

An independent set (IS) in a graph is a set of vertices no two of which are adjacent. An independent set is called maximal (MIS) if there is no independent set that it contains properly. Hopfield network whose stable states are exactly maximal independent sets was developed by Shrivastava [36]. An independent set in a graph is a clique in the complement graph and vice versa. Thus, cliques can be used to find or to enumerate MISs [20, 21]. Finding independent sets (or cliques) has applications in various fields [37]. Combinatorial nets can be used to compute maximal independent sets of graphs in a *distributed self-organization* fashion. A stable states of *a network of bi-stable switches derived from a graph* are exactly maximal independent sets of its underlying graph. Now we consider simple graph *H*, that is, graph without directed, multiple, or weighted edges, and without self loops. Let *C*(*H*) denote the graph obtained from *H* by deleting an each undirected edge (*u*, *υ*) of *H* and adding instead of this edge new two directed edges (*u*, *υ*) and (*υ*, *u*). The set of vertices of *C*(*H*) let be the same as the set of vertices of *H*. Let −1 be a weight of each edge of *C*(*H*). The Bi-stable Switch can be seen as combinatorial net derived from simple graph of order 2.

For example Figure 4b demonstrates the network derived from the graph shown in Figure 4a. The first switch is formed by the subgraph induced by the {1,2} set of vertices of the C(H) network. The second switch is formed by the {2,3} set of vertices. The vertex 2 is common one of these switches, therefore they interact by means of this vertex. Each edge of an underlying graph correspond to a switch in a derived network. If two incident edges share a common vertex then the corresponding switches interact because this vertex have the same state in both of them. We refer to C(H) as the *derived network* of bi-stable switches, and we refer to H as *the underlying graph*.

**Figure 4:**
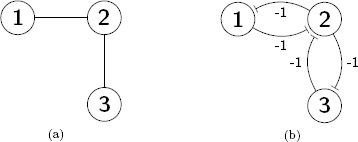
The network C(H) derived from a graph H: (a) The graph H; (b) The network C(H) derived from the graph H;

#### Lemma 1.

*A steady state map M of a combinatorial net that derived from a simple graph does not have two adjacent active vertices u and υ*.

**Proof**. *Assume by contradiction that u and v are adjacent active vertices, then I*(*u*, *M*) < 0 *and the forced state of the u is inactive. If a state is steady and the forced state is inactive then the state of the vertex is inactive, whereas by conditions of the lemma u is active*.

#### Lemma 2.

*If a map M of a combinatorial net that derived from a simple graph is steady and some vertex is inactive then there is at least one active vertex adjacent to it*.

**Proof**. *Assume by contradiction, that there are no active adjacent vertices of *υ*, then influence I*(*υ*, *M*) = 0 *and then forced state of *υ* is active. Hence the map M is steady state, forced state of vertex is equal to current state, then state of the vertex is active in contradiction to conditions of the lemma*.

#### Lemma 3.

*Under steady state map M of the combinatorial net C*(*H*) *derived from simple graph H the set of all active vertices is maximal independent set of H*.

**Proof**. *Let us prove the independence first. Assume by contradiction that the set of active vertices of some steady state map C*(*H*) *is not an independent set of vertices of the graph H. Then there is a pair of adjacent vertices in H, which are simultaneously active under this steady state map M. But by lemma 1 there are no two adjacent active vertices under steady state map. This contradiction proves the independence of the set of active vertices under the steady state map. Now we consider maximality of the set of active vertices under a steady state map. Assume that there is a some inactive vertex which not adjacent to any active vertex. So that it may be added to this set to form the bigger independent set. But if such vertex exist then the map is not steady by lemma 2*.

#### Lemma 4.

*Any maximal independent set of some simple graph H forms the full set of active vertices of some steady state map of the combinatorial net that derived from the H*.

**Proof**. *The desired map M is constructed as follows. Let all vertices from the maximal independent set be active under M, but another vertices let be inactive under M. Evidently, this map M is the steady state map*.

**Theorem about steady states of the network of bi-stable switches**. *Let C*(*H*) *be the combinatorial net derived from simple graph H, then the map M is steady state if and only if the set of all active vertices under map M is maximal independent set of vertices of the graph H*.

**Proof**. *Lemmas 3 and 4 prove the theorem*.

## 6 Conclusions

A similar approach to construct Markov chains for interaction graphs was developed in our earlier works for neural and gene regulatory networks [38, 39, 40, 41, 42]. Both approaches can be used to construct Markov chains of gene regulatory networks. Systems of mutually repressive elements are ubiquitous in nature. The network of bi-stable switches can be used to create models of their stable states and the self-evolution of such systems toward a stable states.

